# Vision rivals audition in alerting humans for fast action

**DOI:** 10.1101/2023.06.02.543380

**Authors:** Niklas Dietze, Christian H. Poth

## Abstract

Successful behaviour requires that humans act promptly upon the ubiquitous rapid changes in the environment. Prompt actions are supported by phasic alertness: the increased readiness for perception and action elicited by warning stimuli (*alerting cues*). Audition is assumed to induce phasic alertness for action faster and more strongly than other senses. Here, we show that vision can be equally effective as audition. We investigated the temporal evolution and the effectiveness of visual and auditory alerting for action in a speeded choice task, while controlling for basic sensitivity differences between the modalities that are unrelated to action control (by matching auditory and visual stimuli according to reaction times in a prior simple detection task). Results revealed that alerting sped up responses, but this happened equally fast and equally strong for visual and auditory alerting cues. Thus, these findings argue that vision rivals audition in phasic alerting for prompt actions, and suggest that the underlying mechanisms work across both modalities.

## 1. Introduction

Humans must rapidly perceive and act upon sudden changes in the environment. For instance, drivers have to slam on the brake when a pedestrian or deer suddenly cuts the road. In such situations, split seconds of reaction time can be vital for survival. To handle time-sensitive situations, the human brain is equipped with powerful alerting mechanisms that support fast action upon stimuli in the environment (Hackley, 2009; Petersen & Posner, 2012). *Alerting effects* are the behavioural signature of these mechanisms: Responses in visual detection and discrimination tasks are faster (Dietze & Poth, 2022; Fan et al., 2002; Hackley, 2009; Poth, 2020), more accurate (Matthias et al., 2010; Petersen et al., 2017; Wiegand et al., 2017), or faster at the cost of errors (Han & Proctor, 2022; McCormick et al., 2019; Posner et al., 1973) when target stimuli are preceded by warning stimuli (alerting cues). Alerting effects are assumed to arise because alerting cues induce phasic alertness, a temporary state of arousal that heightens the overall readiness for perception and action (Posner & Petersen, 1990; Sturm & Willmes, 2001). Arguing for such a global, unspecific modulation of perception and action, the performance benefits occur even though alerting cues provide no information about what response would be correct for the impending target. Thus, rather than promoting a specific modulation through low-level influences on the detection of target stimuli, alerting seems to also support the higher-level processes, such as decision-making for selecting among response alternatives (Fan et al., 2002; Hackley, 2009; Poth, 2020) and perceptual encoding for object recognition (Haupt et al., 2019; Petersen et al., 2017; Wiegand et al., 2017). In line with these global benefits for cognitive processing, alerting can temporarily restore spatially impaired visual perception in neurological patients (Robertson et al., 1998) and alerting trainings can accelerate visual perception in aging (Penning et al., 2021). Alongside clinical applications, the beneficial effects of alerting cues on action in safety-critical situations have long been recognised and used to guide the design of warning signals (Wogalter & Mayhorn, 2005).

Audition elicits the fastest and strongest alerting effects in humans (Bertelson & Tisseyr, 1969; Davis & Green, 1969; Harvey, 1980) and non-human primates (Chapman et al., 1986; Spidalieri et al., 1983). It has been proposed that auditory stimuli are more arousing than stimuli in other modalities such as vision (Nissen, 1977; Ulrich & Mattes, 1996). Across visual tasks, reactions have been found sped up, when auditory alerting cues preceded visual targets shortly (< 150 ms; Posner et al., 1976), but the effect dissipated for longer cue-target onset asynchronies (CTOAs). The effect seems to be phasic and automatic, as it arises quickly but cannot be retained to prolong the readiness for perception and action (Posner et al., 1976). In contrast, this pattern was not observed for visual alerting cues, which has been taken as evidence that the visual modality cannot deliver the fast, short-lasting, and automatic alerting effect. At least for longer CTOAs (≥ 150 ms), it was found that auditory alerting cues and visual alerting cues can induce similar reaction time benefits when controlled for location cueing effects caused by different target and warning signal locations (Rodway, 2005). This shows that the task paradigm and temporal dynamics play a crucial role in alerting effects.

Audition has higher temporal resolution than vision (Artieda & Pastor, 1996), so that auditory alerting cues should affect performance across CTOAs in a temporally more precise way. Across modalities it has been shown, that reaction times generally decrease with increasing CTOAs (Näätänen, 1971). This has been interpreted as evidence that participants’ temporal expectation for the target increases over time, because all used CTOAs were equally likely, so that with every passing CTOA, targets became more and more likely to appear (Niemi & Näätänen, 1981). Temporal expectation contributes to enhanced response readiness (Nobre & van Ede, 2017), and it can support action in addition or interaction to automatic alerting (Lu et al., 2014). The classic studies that compared vision and audition in alerting used fixed or uniformly distributed CTOAs (Bertelson & Tisseyr, 1969; Davis & Green, 1969). Thus, differences in the alerting capability could stem from interactions between their intrinsic temporal dynamics and mechanisms for temporal expectation. Therefore, temporal expectation has to be taken into account when visual and auditory alerting are compared in their effectiveness. It is assumed that this can be achieved by drawing CTOAs from non-aging probability distributions, in which the probability that a target appears now given it has not appeared yet is constant over time (Näätänen, 1971). Such a manipulation has been shown to keep the temporal expectation of the targets constant, and thus reduces its confound with response readiness (Petersen et al., 2017; Weinbach & Henik, 2012). Crucially, it is unknown how differences in auditory and visual alerting effects unfold over longer time-periods using non-aging probability distributions.

The advantage of audition over vision does not necessarily stem from privileged access of audition to mechanisms preparing for perception and action, but could rather be a by-product of audition’s low-level characteristics. That is, reaction times have been found to decline with increasing psychophysical intensity of alerting cues (Adams & Behar, 1966) and auditory processing is generally faster than visual processing (Grondin, 2010; Hove et al., 2013), as transduction times are higher in the auditory than the visual system (Recanzone, 2009; Torre et al., 1995). All this hints at that audition dominates vision in alerting for action due to differences in their low-level characteristics. Arguing against this hypothesis, however, are findings tying audition more closely to mechanisms for action control. Specifically, auditory stimuli seem to result in greater activations in the motor cortex compared with visual stimuli (Fujioka et al., 2012; Grahn & Rowe, 2009). This larger neuronal activity could ultimately translate to faster responses, meaning that audition indeed has a privileged access to alerting for action. For the discussed reasons, the question if audition indeed dominates vision in phasic alertness is still wide open. The privilege of audition for alerting could be inherited from audition’s higher low-level sensitivity and processing speed.

Besides the behavioural effects, differences in alerting can be investigated through physiological markers: Alertness is assumed to be regulated by neuromodulation of cortical processing from the locus coeruleus-norepinephrine system (Aston-Jones & Cohen, 2005; Petersen & Posner, 2012; Posner & Petersen, 1990). Bursts of norepinephrine impact the arousal level and thereby support a wide range of cognitive functions. This neuronal activation can be indirectly assessed with changes in pupil size (Nieuwenhuis et al., 2011; Reimer et al., 2016). The size of the pupil increases with the intensity of an alerting cue (Petersen et al., 2017), in line with the idea that it reflected intensity-dependent arousal. Thus, assuming that this pupil dilation was a measure for arousal (e.g., Aston-Jones & Cohen, 2005; Joshi & Gold, 2020; Poth, 2021), the advantages of audition for eliciting phasic alertness due to increased arousal should become manifest in stronger pupil dilations for auditory alerting cues than for cues in other modalities. In addition to pupil responses, saccade latencies have also been linked to the current state of alertness (Bocca & Denise, 2006). It has been found that saccadic latencies are facilitated when preceded by an alerting cue (Ross & Ross, 1980). For these reasons, not only reaction times but also physiological arousal measures should be included in comparisons between alerting modalities.

Therefore, to settle this issue, we investigated the effectiveness and temporal evolution of visual and auditory alerting in a choice reaction task, while equating for the simple detection times between the modalities. First, participants performed a simple detection task to identify the intensities of auditory and visual alerting cues that yielded to similar reaction times (Experiment 1). Following the matching procedure, participants performed a speeded choice with the individually determined intensities of visual and auditory alerting cues (Experiment 2). However, unlike matching procedures focusing on perceived stimulus intensities (cf. Stevens & Marks, 1965), we explicitly did not equalise the stimulus intensities at a perceptual level but regarding speeded responses. Using reaction times, we controlled for the speed advantage of audition with respect to earlier processing that is required for a simple reaction without a choice. In contrast to simple reactions, choice reactions measure additional processes of action control, such as response selection and error avoidance (Stuss et al., 2005). To control for temporal expectation, CTOAs across trials were drawn from a non-aging probability distribution, following the gold standard outlined above (Weinbach & Henik, 2012). Surprisingly, however, we still found traces of temporal expectation in our results, casting doubt on a clear dissociation of temporal expectation and alerting. Our findings show that with matched alerting cues based on the known reaction times in a simple detection task, visual alerting is just as effective as auditory alerting, both in terms of behavioural performance as well as physiological arousal responses (pupil dilation and latencies of spontaneous saccades). As such, the findings show that vision can rival audition when the stimulus intensities are matched based on the simple detection times of visual and auditory alerting cues. Phasic alertness seems to arise equally from different senses, which may hint at modality-general alerting mechanisms.

## 2. Method

### 2.1. Participants

Seven participants from the university including one of the authors (participant ND) took part in the preregistered experiments (https://osf.io/fud4k). They were between 21 and 41 years old (*median* = 27 years), six were female and one was male. Due to the psychophysical nature of the present investigation and the large number of trials (7000 trials in Experiment 1, 52500 trials in Experiment 2), we limited the sample size to seven participants (one more participant than initially preregistered because one of the authors participated in the study) which should suffice to establish the existence of effects in the population as long as all participants show qualitatively similar results (Anderson & Vingrys, 2001; Smith & Little, 2018). All participants reported normal or corrected-to-normal vision, normal hearing, and gave written informed consent before participation. The study was in accordance with the ethical guidelines of the German Psychological Association (DGP) and approved by university’s ethics committee.

### 2.2. Apparatus and stimuli

Both experiments took place in the same dark room with the only light sources being the display monitor and the operator monitor located behind the participant. Participants were seated in front of the preheated display monitor (Poth & Horstmann, 2017) at a viewing distance of 71 cm with their head on a chin-and-forehead rest. Eye-movements were recorded with an Eyelink 1000 (SR Research, Ottawa, ON, Canada) at a sampling rate of 1000 Hz. The CRT monitor (G90fB, ViewSonic, Brea, CA, USA) ran at a refresh rate of 85 Hz and a resolution of 1024 x 768. Stimuli were controlled using MATLAB R2014b (The MathWorks, Natick, MA, USA) and the Psychophysics toolbox (Brainard, 1997; Kleiner et al., 2007; Pelli, 1997). Responses were collected with a standard external computer mouse placed in front of the participants. Auditory stimuli were presented using loudspeakers placed 45 cm to the left and right of screen centre (Philips Multimedia Speaker System A 1.2 Fun Power/MMS 101, Philips, Amsterdam). All visual stimuli were white figures (102.6 cd/m^2^) presented on a black background (0.03 cd/m^2^). A filled circle with a diameter of 0.28° of visual angle was used as a fixation point for both experiments. In Experiment 1, the target stimulus was either a larger frame with a diameter of 5° of visual angle presented at the centre of the screen or a sine tone with a frequency of 700 or 900 Hz (both frequencies occurring equally often and in random order across trials) to avoid habituation effects. In Experiment 2, the same stimuli were used as alerting stimuli. The target stimulus was a filled square of 0.5° of visual angle which appeared either 6.5° to the left or to the right of screen centre. To check the precise timing of the custom experimental program, we measured the auditory and visual stimuli externally (cf. Poth et al., 2018) with a microphone capsule and a BPW-34 photodiode, sampled at 2.5 kHz using a TDS 2022 B oscilloscope (Tektronix, Beaverton, OR, USA). Fifteen runs, in which an auditory alerting stimulus or a visual alerting stimulus preceding a visual target stimulus with CTOAs from 153 ms to 529 ms were measured. We found that the CTOAs for the visual stimuli were on average 0.2 ms shorter than programmed. For the runs with both modalities, the CTOAs were on average 1.5 ms shorter.

### 2.3. Procedure of Experiment 1

The first experiment served as a baseline measurement to match visual and auditory cues based on the trimmed mean reaction times. At the beginning of the experiment and after 250 trials or 20 broken fixations, a 9-point calibration of the eye-tracker was conducted. In this experiment, 10 logarithmically spaced stimulus intensities for both modalities were presented each 50 times randomly throughout the session after a random fixation period of 750 ms to 1250 ms drawn from a uniform distribution for a duration of 50 ms. Participants had to respond as quickly as possible to the stimulus onset. An additional 1500 ms for pupillometry followed after response collection. Visual intensity varied from 5.79 cd/m² to 102.6 cd/m² and auditory intensity varied from 36.6 db(A) to 64.2 db(A). Each participant conducted one session (1000 trials for each participant, 7 x 1000 = 7000 trials for the whole sample). On average, the time to complete a session was about 65 min.

### 2.4. Procedure of Experiment 2

In Experiment 2, the matched stimuli based on the trimmed mean reaction times in the prior simple detection task for each participant were used to measure the alerting effects for both modalities. The same calibration conventions as in Experiment 1 were applied to Experiment 2. Trials started with the presentation of the fixation point. After a random fixation period of 750 ms to 1250 ms drawn from a uniform distribution, either an auditory cue (1/3 of trials), a visual cue (1/3 of trials) or no cue (1/3 of trials) was presented for 50 ms. The CTOAs were drawn from a non-aging probability distribution in steps of 47.04 ms from 153 ms to 3965 ms with a probability of 0.1. After the waiting period, the target stimulus was presented for 200 ms. If participants moved their eyes outside of the fixation window of 2.5° before target onset, the trial was randomly repeated during the remaining sequence. Participants were asked to respond as quickly as possible to the location of the (6.5° to the left or to the right of screen centre) target by pressing the corresponding mouse button (Fig. 1). Again, an additional 1500 ms for pupillometry followed after response collection. Each participant conducted 10 sessions (10 x 750 = 7500 trials for each participant, 7 x 10 x 750 = 52500 trials for the whole sample). However, technical problems prevented recording of 268 trials resulting to a final sample of 52232 trials. Controlling for fatigue, no more than two sessions were completed on the same day. Between two sessions a maximum of seven days elapsed. All sessions were completed on average within a period of six weeks (*min* = 14 days, *max* = 94 days). The time to complete a session was about 60 min. Before and after each session, participants filled out a short questionnaire on subjective relaxation as part of a larger assessment across studies (cf. Steghaus & Poth, 2022).

### 2.5. Behavioural analyses

Statistical analyses were performed in R (4.0.5, R Core Team, 2021). Following the preregistered protocol, reaction times for each experimental condition were compared using a Bayesian multilevel model created in the Stan computational framework (https://mc-stan.org) accessed with the R-package brms (2.16.3, Bürkner, 2017). We used a Bayesian model instead of a frequentist approach to quantify evidence in favour of the null hypothesis. In addition to several other advantages, Bayesian models are more robust with small sample sizes (Stegmueller, 2013). We analysed reaction times as a function of alerting condition and CTOA with a random intercept per participant, a random slope per alerting condition and a random slope per CTOA. The model fit was assessed with the expected log predictive density based on approximate leave-one-out cross-validations. For both fixed effects and the random effects of the final multilevel model, we specified conservative priors. The model was run with 4 MCMC chains and 4000 iterations for each chain. Complementary, pairwise comparisons between the alerting effects (mean difference between no cue trials and alert trials) across all CTOAs and participants were tested using paired *t*-tests with Cohen’s *d_z_* effect size (Cohen, 1988) and Bayesian *t*-tests using standard settings of the R-package BayesFactor (0.9.12-42; Morey & Rouder, 2021), whose Bayes factors quantify evidence in favour of the null hypothesis (*BF_01_*) or evidence in favour of the alternative hypothesis (*BF_10_*). All data processing and visualisations were made using the tidyverse libraries (Wickham et al., 2019) and data.table (Dowle et al., 2023). Error trials (1.5 %), trials with CTOAs greater than 1518 ms (3.9 %) and trials for which the reaction time was greater than 2.5 SD (2.3 %) were removed from the analyses.

### 2.7. Pupil analyses

The average pupil size after the pupil normalised from the pupillary light reflex was computed and compared between conditions. Baseline pupil diameter on each trial was defined as the mean pupil diameter in the interval 100 ms prior to cue onset. We used the subtraction method to compute baseline corrected pupil responses for each participant. Mean alerting induced pupil dilations (mean difference between no cue trials and alert trials) for each participant were compared using paired *t*-tests with Cohen’s *d_z_* effect size and Bayesian *t*-tests. The same exclusion criteria as the behavioural analyses were applied to the pupil data.

### 1.1. Saccade analyses

Mean saccade latencies of the first saccade relative to target onset were extracted and compared between alerting conditions (mean difference between no cue trials and alert trials). We also used paired *t*-tests with Cohen’s *d_z_* effect size and Bayesian *t*-tests. On top of the previous exclusion criteria, trials were not used if the latency of the first saccade was less than 0 ms or greater than 1000 ms (15.8 %) relative to target onset.

## 3. Results

### 1.1. Behavioural results

The matched intensities of Experiment 1 are presented in Table 1 and the mean reaction time results of Experiment 2 are visualised in Figure 2. In Experiment 2, reaction times were generally shorter when preceded by a visual alerting, *t*(209) = 18.490, *p* < .001, *d_z_* = 1.276, *BF_10_* = 2.9 x 10^42^, and an auditory alerting cue, *t*(209) = 17.434, *p* < .001, *d_z_* = 1.203, *BF_10_*= 1.8 x 10^39^, compared to the no cue condition (cf. Fan et al., 2002; Hackley, 2009; Poth, 2020). Differences in reaction times between trials with an alerting cue and trials without an alerting cue diminished with increasing CTOAs. In the early CTOAs, the phasic alerting effect was about 30 ms. In later CTOAs, it decreased up to a point where differences between conditions were no longer present. This was particularly due to the change in the no cue condition. Reaction times decreased with longer CTOAs which is surprising considering the use of non-aging probability distributions to control for temporal expectation. Thus, it seems that traces of temporal expectation contributed to the present results. We also found that visual alerting was not inferior to auditory alerting. Based on the observations of the posterior distribution, auditory alerting was with 98.5 % confidence not better than visual alerting. This means that the hypothesis being tested (auditory alerting > visual alerting) was in 98.5 % of the posterior samples more likely than the alternative hypothesis (auditory alerting ≤ visual alerting). The posterior mean of the difference was −3.3 ms with a 95 % credible interval from −6.2 ms to −0.4 ms. On average participants responses were 24.2 ms (*SD* = 19.0 ms) faster following the visual alerting cue and 20.9 ms (*SD* = 17.4 ms) faster following the auditory alerting cue (Fig. 3). In summary, near 100 % of the tested cases revealed that auditory alerting was not superior to visual alerting. This was supported by the pairwise comparisons, *t*(209) = −4.319, *p* = 1.000, *d_z_* = −0.298, *BF_01_* = 73217, showing that reaction times following an auditory alerting cue were not shorter than reaction times following a visual alerting cue. Importantly, these findings cannot be explained by a speed-accuracy trade-off as the overall error rate was low (1.5 %) and differences between the cue conditions only very minor (no cue: 0.4 %, visual: 0.6 %, auditory cue: 0.5 %). Pairwise comparisons showed no differences between the no cue and visual alerting cue, *t*(209) = −0.292, *p* = .771, *d_z_* = 0.020, *BF_01_* = 12.429, no cue and auditory alerting cue, *t*(209) = −0.476, *p* = .635, *d_z_* = 0.033, *BF_01_* = 11.593, and visual alerting cue and auditory alerting cue, *t*(209) = 0.215, *p* = .830, *d_z_* = 0.015, *BF_01_* = 12.671. Note, that these findings did not change with the exclusion of participant ND. The same analyses without participant ND are reported in the supplementary material.

**Figure 2.**
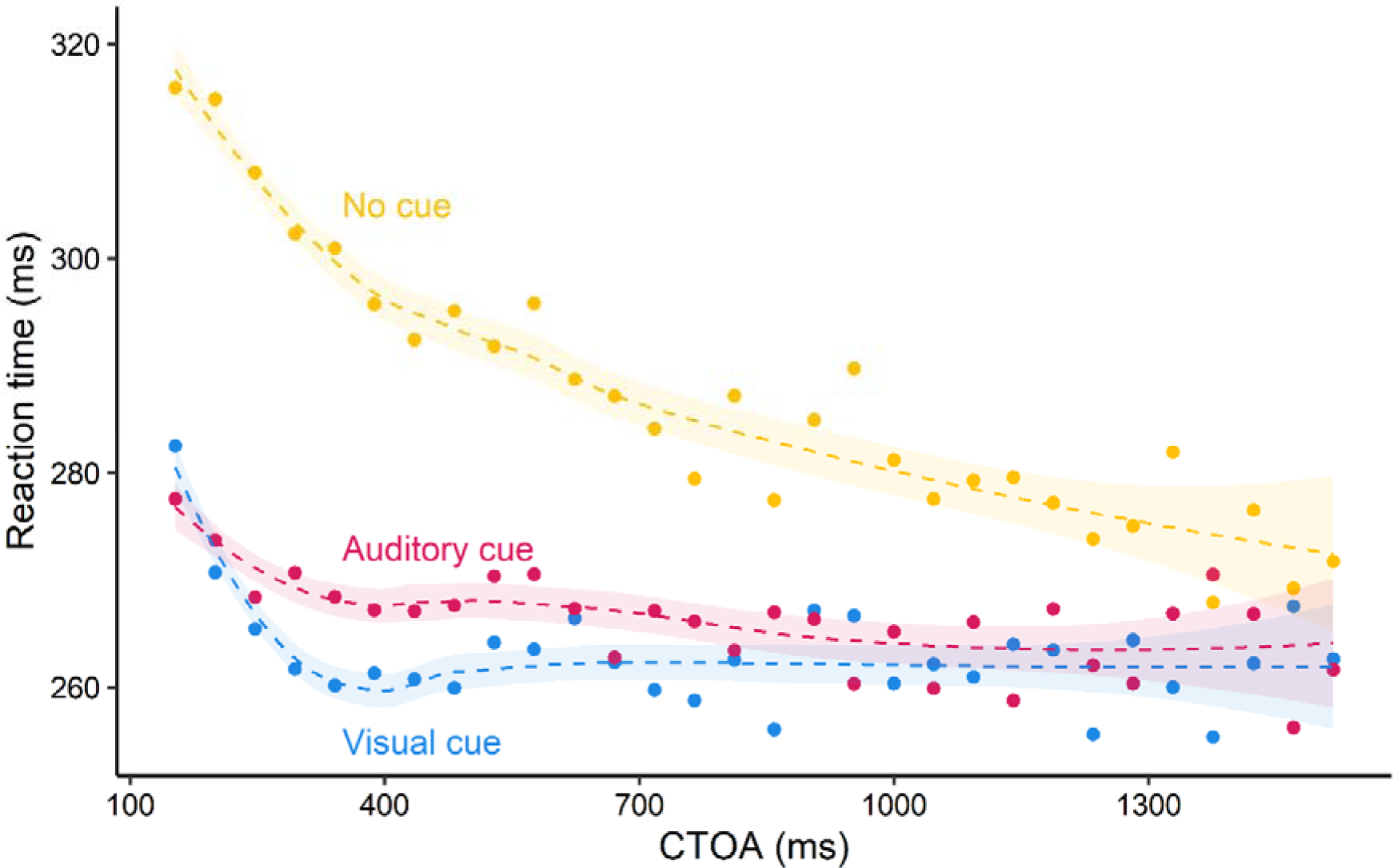
Manual responses of Experiment 2. Participants’ mean reaction times fitted with a polynomial regression in the three experimental conditions. Confidence bands depict the 95 % confidence interval.

**Table 1.** Matched stimulus intensities for each participant based on the mean reaction times.

| Participant | Auditory intensity<br>Decibel (A) | RT (ms)<br>Mean (SD) | Visual intensity<br>Candela (cd/m<sup>2</sup>) | RT (ms)<br>Mean (SD) |
| --- | --- | --- | --- | --- |
| 1 | 36.6 | 206 (31) | 8.11 | 208 (33) |
| 2 | 52.3 | 272 (77) | 55.56 | 265 (51) |
| 3 | 36.6 | 280 (90) | 11.27 | 273 (88) |
| 4 | 36.6 | 259 (73) | 15.60 | 271 (74) |
| 5 | 36.6 | 340 (71) | 11.27 | 351 (72) |
| 6 | 36.6 | 261 (80) | 11.27 | 266 (74) |
| 7 | 64.2 | 307 (15) | 102.6 | 300 (84) |

### 1.2. Pupillary results

Pupil responses were observed in all three cue conditions (Fig. 4 a). As such, part of the responses must be attributed to target onset and detection. Nevertheless, differences between conditions provide an estimate about the arousal induced by the alerting signals (Petersen et al., 2017). Directional pairwise comparisons between the pupil alerting effects, *t*(209) = 0.381, *p* = .352, *d_z_* = 0.026, *BF_01_* = 0.545, revealed indecisive evidence on the hypothesis that auditory alerting is superior to visual alerting. However, pairwise comparisons testing the non-directional hypothesis, *t*(209) = 0.381, *p* = .704, *d_z_* = 0.026, *BF_01_*= 12.068, showed that the pupil dilations were not different between modalities. On average pupil responses from baseline for the interval from 1500 ms to 1600 ms relative to cue onset were .053 mm *(SD* = .123 mm) following the visual alerting cue and .052 mm *(SD* = .115 mm) following the auditory alerting cue.

**Figure 4.**
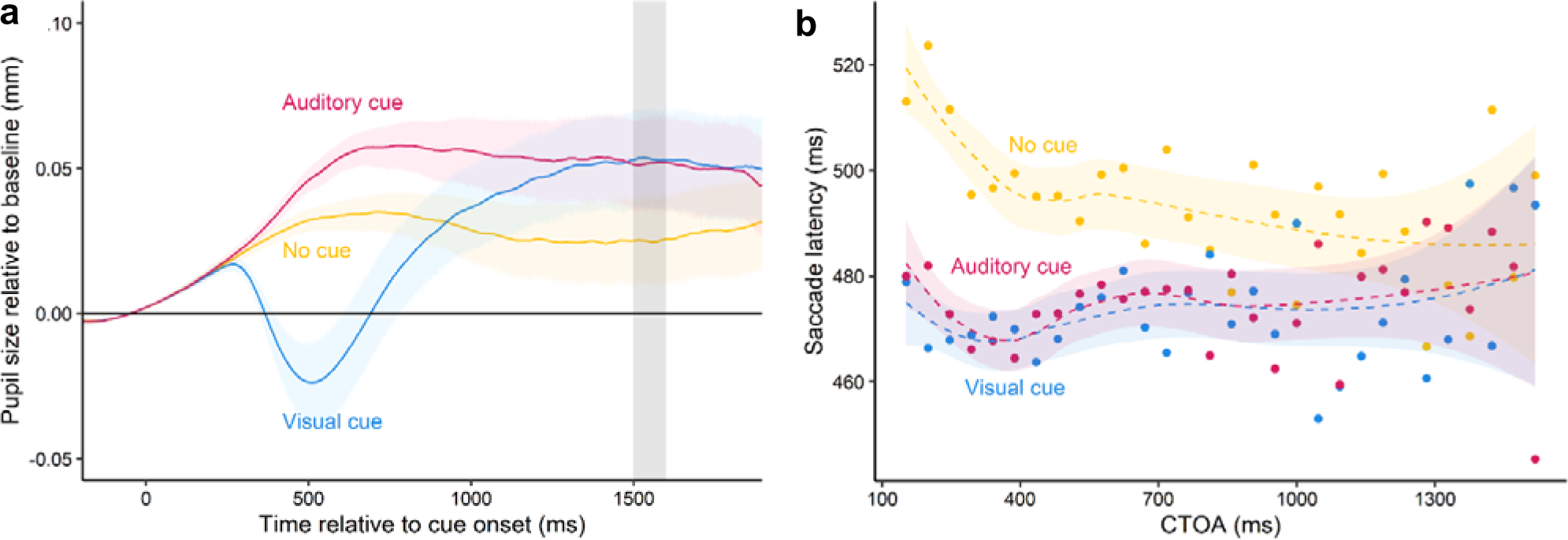
Pupil responses and saccadic latencies of Experiment 2. **a** Grand averages of pupil dilation for the three experimental conditions relative to cue onset. Shaded area shows the analysed time period after the pupil recovered from the pupillary light reflex. Confidence bands depict the 95 % confidence interval. **b** Saccadic latencies fitted with a polynomial regression in the three experimental conditions. Confidence bands depict the

### 1.1. Saccadic latency results

As an additional eye-movement marker for phasic alertness, we investigated the latencies of the oculomotor response. Similar to the manual reaction times, saccadic latencies after target onset were shortened when preceded by a visual alerting cue, *t*(209) = 6.501, *p* < .001, *d_z_* = 0.449, *BF_10_* = 1.4 x 10^7^, and an auditory alerting cue, *t*(209) = 7.467, *p* < .001, *d_z_* = 0.515, *BF_10_* = 3.1 x 10^9^, compared to the no cue condition (Fig. 4 b) (cf. Ross & Ross, 1980). Again, using directional pairwise comparisons between the saccadic alerting effects, *t*(209) = 0.124, *p* = 0.451, *d_z_* = 0.009, *BF_01_* = 0.822, revealed indecisive evidence on the hypothesis that auditory alerting leads to faster saccade latencies than visual alerting. But, pairwise comparisons testing the non-directional hypothesis, *t*(209) = 0.124, *p* = 0.902, *d_z_*= 0.009, *BF_01_*= 12.865, showed that the saccadic alerting effects are not different between modalities. On average saccadic latencies were 18.7 ms (*SD* = 41.8 ms) faster following a visual alerting cue and 19.1 ms (*SD* = 37.0 ms) faster following an auditory alerting cue compared to the condition without a preceding cue. Overall, the saccade responses occurred on average 205 ms after the motoric response (Fig. 5 a), which suggests that the motoric response delayed the temporal execution of an eye-movement (Reimer et al., 2020). Note, however, most of the spontaneous saccades landed close to screen centre (Fig. 5 b). Only 1.6 % of the saccades were directed towards the target locations (no cue: 0.5 %, visual: 0.5 %, auditory cue: 0.6 %). Most importantly, across all saccades no differences in the horizontal landing positions were found between the no cue and visual alerting cue, *t*(209) = −1.657, *p* = .099, *d_z_* = −0.114, *BF_01_* = 3.375, no cue and auditory alerting cue, *t*(209) = −1.947, *p* = .053, *d_z_* = −0.134, *BF_01_* = 2.032, and visual alerting cue and auditory alerting cue, *t*(209) = 0.021, *p* = .984, *d_z_* = 0.001, *BF_01_* = 12.960. Thus, although participants’ saccades were facilitated, we did not find that visual or auditory alerting promoted orienting towards the target.

**Figure 5.**
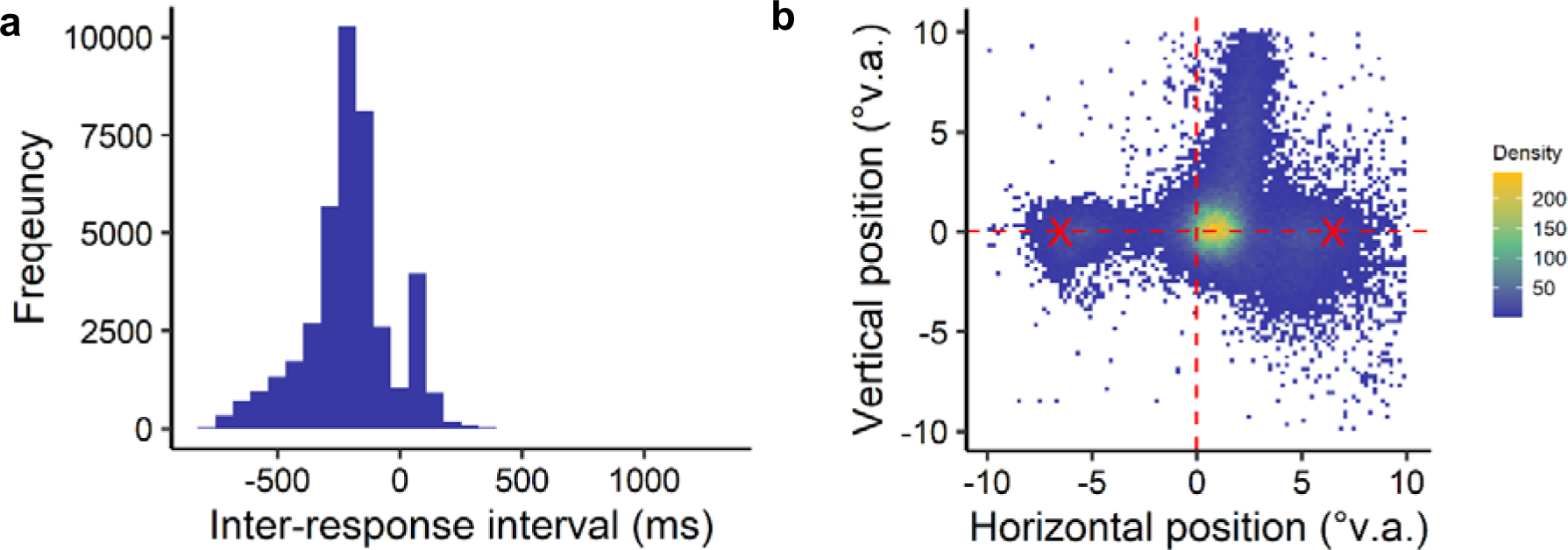
Inter-response intervals and saccadic landing positions of Experiment 2. **a** Frequency distribution of the inter-response interval, the differences between reaction times and saccade latencies. Negative inter-response intervals indicate that the manual responses were executed prior to the saccades. **b** Density plot of average eye position after the first saccade in degrees of visual angle. Bright yellow squares show the locations with the highest density of saccade landing positions. Red X’s depict the two possible

## 4. Discussion

We discovered that in contrast to common views, visual and auditory alerting supports fast action equally well. We employed a novel procedure for matching intensities of alerting cues according to reaction times in a prior simple detection task to control for low-level differences between audition and vision. For alerting cues with matched intensities, auditory and visual alerting both elicited faster responses, and this was the case across CTOAs up to around 1000 ms. This phasic alerting effect was consistent for both modalities across all participants. Crucially, these findings contradict classic studies that have reported large auditory advantages of 31 ms with a foreperiod of 30 ms (Bertelson & Tisseyr, 1969) and 36 ms with a foreperiod of 50 ms (Davis & Green, 1969). In the present study, we used a wide range of CTOAs including very short foreperiods drawn from a non-aging probability distribution. Nevertheless, auditory alerting was never significantly better than visual alerting. In fact, participants responses were on average even slightly faster (for 3.4 ms) with a preceding visual cue. Thus, this indicates that the previously reported advantages of audition for alerting (Bertelson & Tisseyr, 1969; Davis & Green, 1969; Harvey, 1980) could be reflected by the higher sensitivity and processing speed of audition. Once controlled for these low-level characteristics of audition, vision rivals audition in alerting humans for perception and action.

Increasing the level of alertness is also expressed in physiological arousal responses. For instance, the pupil dilates once participants are alerted (Aston-Jones & Cohen, 2005; Petersen et al., 2017), and the magnitude of dilation seems positively correlated with intensity levels of the alerting cue (Petersen et al., 2017). Thus, for the present study, differences in pupil responses would highlight any modality specific influences. However, we observed similar pupil responses between the auditory and visual alerting modality. Although at the individual level, pupil responses varied due to differences in intensity levels, dilations in the no cue condition were always systematically weaker than the alerting conditions. In addition, we found a similar relationship in saccadic latencies. Saccadic latencies were equally facilitated when preceded by auditory and visual alerting cues. In contrast to manual reaction times, spontaneous saccades are more automatic and less prone to confounding factors such as the speed-accuracy trade-off and response criterion (Ross & Ross, 1980). Therefore, these patterns of results offer converging evidence that there is no auditory dominance in phasic alertness.

In the present study, we focused on visual targets for two main reasons. First, studies that have reported an auditory dominance found the greatest effects on behaviour with auditory stimuli preceding visual targets (Bertelson & Tisseyr, 1969; Davis & Green, 1969; Harvey, 1980). Second, the visual domain is most relevant for action control (Posner et al., 1976). Previous accounts have postulated that cross-modal auditory advantages have been caused by different locations for visual alerting cues and visual targets (Rodway, 2005; Turatto et al., 2002). In classic experiments, visual cues typically appear at different locations than the visual target while auditory alerting cues only convey little spatial information (Bertelson & Tisseyr, 1969; Davis & Green, 1969; Harvey, 1980; Posner et al., 1976). Hence, it is easier to shift attention from an auditory cue to a visual target than from a visual cue to a visual target (Rodway, 2005). This is in line with the hypothesis that visual input typically engages more attention than any other modality (Posner et al., 1976). Since attention is highly focused to the visually cued location, it makes it more difficult to shift attention away to the following visual target (Posner et al., 1976). One way to minimise this location cueing effect, would be to use cues that do not convey any spatial properties (Wright & Richard, 2003). For our present study we used a relatively large white frame presented around screen centre which should not have caused any location specific sensory processing (Matthias et al., 2010). Even when the present manipulation did not fully control for such an influence, our results argue against a cost induced by an additional attentional shift caused by the visual warning signal. That is, we did not find any auditory advantage in the early CTOAs. However, the present data cannot rule out any differences in CTOAs below 153 ms.

We found that the magnitude of the phasic alerting effect decreased with longer CTOAs. In particular, it was observed that the mean reaction times in the condition without a cue converged in an exponential fashion with the other two alerting conditions at around 500 ms. This contradicts with findings suggesting no changes in phasic alertness with CTOAs up to 1000 ms (Lu et al., 2014). Moreover, it has been reported when using non-aging probability distributions that reaction times slightly increase as a function of CTOA (Lu et al., 2014; Saban et al., 2019). Typically, faster reaction times with increased CTOAs are only observed when using aging or accelerated-aging probability distributions (Fuentes & Campoy, 2008; Lu et al., 2014). This is because the probability for the target appearance becomes greater over time, which in turn leads to a build-up in temporal expectation (Coull, 2009). The present pattern of results was quite similar to previous studies that did not use non-aging probability distributions, and thus questions the procedure to control for temporal expectation. Here, we used a fine-grained temporal resolution with time-steps of 47 ms and many CTOAs per bin up to 1518 ms which draws a clear picture of the temporal evolution of reaction times. This suggests that response readiness cannot be completely dissociated from temporal expectancy when using non-aging probability distributions as both processes might be triggered by the alerting signal. In fact, more recently it was proposed that that the hazard rate, which is kept constant using non-aging probability distributions, may not be suitable to account for temporal expectancy (see also Grabenhorst et al., 2019).

The present study focused on the most common warning modalities: vision and audition. It would be interesting to find out whether the same findings on alerting extend to other modalities such as touch and temperature. Future studies using the same paradigm would benefit from smaller intensity increments of the alerting cues, so that the detection times could be equated more precisely. In addition, a monitor with ultra-high temporal resolution (Poth et al., 2018) could potentially unravel minor differences in the time-course between the modalities.

In conclusion, we found that alerting works equally well with visual and auditory alerting cues as long as low-level differences are controlled for. This delivers first time evidence that phasic alertness exerts its effects in a modality-general way because the underlying mechanisms do not grant privileged access to a particular sense.

## Statements and Declarations

### Ethics approval

The study was in accordance with the ethical guidelines of the German Psychological Association (DGP) and approved by Bielefeld University’s ethics committee.

### Consent to participate

All participants gave written informed consent before participation.

### Availability of data and materials

All data, analysis code, and experiment code have been made publicly available on the Open Science Framework (https://osf.io/7g4d5/). The design, sampling and analysis plan were preregistered (https://osf.io/fud4k).

### Author Contributions

Conceptualisation: CHP; Methodology: ND, CHP; Software: ND, CHP; Formal analysis: ND, CHP; Investigation: ND, Data Curation: ND, CHP; Writing – Original Draft: ND, CHP; Writing – Review & Editing: CHP, ND; Visualisation: ND; Supervision: CHP; Funding Acquisition: CHP

## Acknowledgements

We acknowledge support for the publication costs by the Open Access Publication Fund of Bielefeld University and the DFG.

## Supplemental material

Using the sample without participant ND (*n* = 6), we also found that visual alerting was not inferior to auditory alerting. Auditory alerting was with 98.2 % confidence not better than visual alerting. This means that the hypothesis being tested (auditory alerting > visual alerting) was in 98.2 % of the posterior samples more likely than the alternative hypothesis (auditory alerting ≤ visual alerting). The posterior mean of the difference was −3.6 ms with a 95 % credible interval from −7.1 ms to −0.3 ms. On average participants responses were 26.6 ms (*SD* = 19.1 ms) faster following the visual alerting cue and 23.0 ms (*SD* = 17.7 ms) faster following the auditory alerting cue (Fig. S1). This was supported by the pairwise comparisons, *t*(179) = −4.151, *p* = 1.000, *d_z_* = −0.309, *BF_01_* = 549, showing that reaction times following an auditory alerting cue were not shorter than reaction times following a visual alerting cue.

---

**Figure S1.**
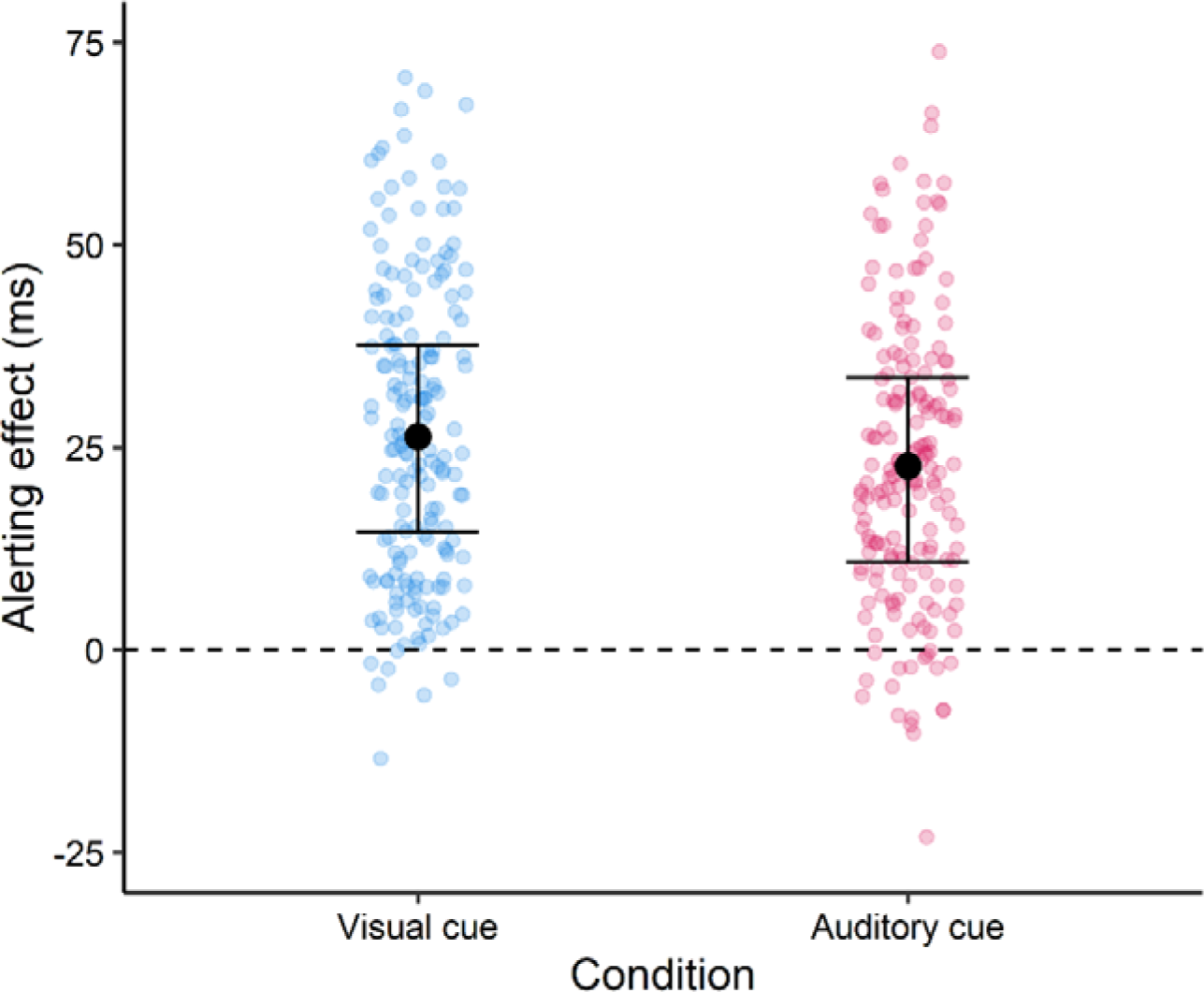
Alerting effects of Experiment 2 without participant ND. Based on the multilevel model, solid red points show the fitted values for the aggregated alerting effects of each participant and CTOA. Error bars depict the 95 % credible

